# Optogenetic urothelial cell stimulation induces bladder contractions and pelvic nerve afferent firing

**DOI:** 10.1101/2023.02.17.528980

**Authors:** Gabriella L Robilotto, Olivia J Yang, Firoj Alom, Richard D Johnson, Aaron D Mickle

## Abstract

Urothelial cells, which play an essential role in the barrier function, are also thought to play a sensory role in bladder physiology by releasing signaling molecules in response to sensory stimuli that act upon adjacent sensory neurons. However, it is challenging to study this communication due to the overlap in receptor expression and proximity of urothelial cells to sensory neurons. To overcome this challenge, we have developed a mouse model where we can directly stimulate urothelial cells using optogenetics. We have crossed a uroplakin II-cre mouse (UPK2-Cre) with a mouse that expresses the light-activated cation channel, Channelrhodopsin-2 (ChR2), in the presence of cre-expression. Optogenetic stimulation of urothelial cells cultured from UPK2-ChR2 initiates cellular depolarization and release of adenosine triphosphate. Cystometry recordings demonstrate that optical stimulation of urothelial cells increases bladder pressure and pelvic nerve activity. Increases in bladder pressure persisted, albeit to a lesser extent, when the bladder was excised in an *in vitro* preparation. The P2X receptor antagonist, PPADS, significantly reduced optically evoked bladder contractions *in vivo* and *ex vivo*. Further, corresponding nerve activity was also inhibited with PPADS. Our data suggest that urothelial cells can initiate robust bladder contractions via sensory nerve signaling or contractions through local signaling mechanisms. This data supports a foundation of literature demonstrating communication between sensory neurons and urothelial cells. Importantly, with further use of these optogenetic tools, we hope to scrutinize this signaling mechanism, its importance for normal micturition and nociception, and how it may be altered in pathophysiologic conditions.

**Animal Studies:** All the procedures involving mice and mouse tissue performed in this study were approved by the University of Florida Institutional Animal Care and Use Committee and in strict accordance with the US National Institute of Health (NIH) Guide for the Care and Use of Laboratory Animals.

**Significance:** It has been appreciated for almost two decades that urothelial cells play a sensory role in bladder function. However, it has been particularly challenging to study this communication as both sensory neurons, and urothelial cells express the same sensory receptors. Here we demonstrate an optogenetic technique to specifically stimulate urothelial cells and evaluate the effects on bladder physiology. We used this technique to show that specific urothelial stimulation resulted in bladder contractions. This approach will have a long-lasting impact on how we study urothelial-to-sensory neuron communication and the changes that occur under disease conditions.

## Introduction

Urothelial cells, the endothelial cells that line the bladder lumen, provide a protective barrier between the urine and the bladder wall. In addition to their barrier function, urothelial cells express sensory receptors and can communicate with adjacent sensory neuron nerve endings^1,2^. Pathological changes to the urothelium, including changes in release factors, have been identified in various bladder diseases, including interstitial cystitis/bladder pain syndrome^3–7^ and overactive bladder^8–11^.

A sensory role of urothelial cells has been postulated for more than two decades, with the primary evidence being 1) they express many sensory receptors, including Piezo^12–14^, ASIC^15,16^, TRPV1^17^, TRPV4^18^, and other TRP^19,20^ channels, 2) they can release several neurotransmitters and signaling peptides (substance P^2,21^, adenosine triphosphate (ATP)^11,14,22–24^, acetylcholine^25,26^, nitric oxide^27,28^, prostaglandins^8–10^) and, 3) they are in proximity to terminations of sensory fibers innervating the urothelial layer^29,30^ (Reviewed in^31,32^). The seminal papers describe a release of ATP following mechanical stimulation of the urothelium^22^, and the sensory defects observed in mice lacking functional P2X_2_ and P2X_3_ receptors^33,34^. Here they showed that stretch-induced and ATP-induced activation of sensory afferents in the bladder were reduced in P2X_2_ and P2X_3_ knock-outs, and ATP release was proportional to sensory nerve activity likely due to urothelial-stretch-induced ATP release activating sensory neurons. However, because of the ubiquitous expression of purinergic receptors in different bladder cell types, including interstitial and urothelial cells, and the various signaling molecules released by urothelial stimulation, more complicated signaling mechanisms cannot be ruled out.

While we know sensory afferents can receive signals from urothelial cells, their exact role in bladder sensation is difficult to delineate. The primary challenge that arises when interrogating these signaling mechanisms is that many urothelial cells and sensory neurons express the same receptors and transmitters, making interpretation of traditional pharmacology difficult. Therefore, understanding the pathophysiology of urothelial cells and visceral sensory neurons, both individually and as a network, is critical for effective therapy development.

To overcome these challenges, we have developed a novel transgenic mouse model that expresses channelrhodopsin-2 (ChR2), an optically activated non-selective cation channel, specifically in the urothelium. With this model, we can utilize optogenetics by activating urothelial-expressed ChR2 to modulate their activity directly. With this technique, we can isolate the effects of urothelial stimulation from other sensory stimuli and study the impact of urothelial stimulation alone.

Here we demonstrate that optogenetic urothelial stimulation alone can evoke bladder contractions of similar magnitude as voiding contractions. Further, the stimulation of urothelial cells results in evoked action potentials in the spinal cord projecting pelvic sensory afferents, and stimulation of these cells in an excised bladder preparation also results in bladder contractions. Both *in vivo* and *ex vivo* contractions partially depended on P2X receptor signaling.

## Materials and Methods

### Animals

All the procedures involving mice and tissue performed in this study were approved by the University of Florida Institutional Animal Care and Use Committee and in strict accordance with the US National Institute of Health (NIH) Guide for the Care and Use of Laboratory Animals. All procedures were performed during the light cycle (0600 – 1800) and were housed on a 12-hour light/dark cycle with access to food and water ad libitum. For our transgenic model, we crossed the Ai32 line that expresses ChR2 (024109, The Jackson Laboratory) with uroplakin II-Cre (029281, The Jackson Laboratory) that expresses UPK2 in the bladder urothelium^35,36^. This novel transgenic line expresses ChR2 within the urothelium, allowing for the targeted activation of urothelial cells with blue light. Both male and female mice were used.

### Immunohistochemistry

Animals were anesthetized with a 1.2 g/kg ketamine-xylazine cocktail and transcardially perfused with 0.01M phosphate-buffered saline (1x PBS). The mice were then perfused with freshly prepared 4% paraformaldehyde (PFA) (19210, Electron Microscopy Sciences) and the bladder was removed and post-fixed in 4% PFA at 4°C overnight. Bladders were then transferred to 30% sucrose in 1x PBS at 4°C. After cryopreservation (∼24 hours), tissues were embedded in Optimal Cutting Temperature Medium (23-730-571, Fisher Scientific) and stored at -80°C.

Bladder tissue was sectioned using cryostat onto microscope slides (12-550-15, Fisher Scientific) at 20 μm. Slides were stored at -20°C until staining. Slides were post-fixed in 4% PFA for 5 minutes followed by 3 washes in 1x PBS for 5 minutes each. Non-specific antibody binding was blocked for 1 hour at room temperature with a blocking solution containing: 5% normal goat serum (006001, SouthernBiotech™), 0.3% Triton™ X-100 (X100, Sigma-Aldrich) in 1x PBS. Tissue sections were incubated with primary antibodies overnight at 4°C, followed by 3 washes in 1x PBS for 5 minutes each (see **Table 1** for antibody details). Thereafter, sections were incubated with secondary antibodies for 45 minutes at room temperature. All antibodies were diluted in blocking solution. Slides were mounted using VECTASHIELD Vibrance Mounting Media (H-1700-10, Vector Labs). All images were captured with the Keyence BZ-X Series all-in-one fluorescence microscope and BZ-X Viewer, and processed with Fiji ImageJ (1.53t, NIH).

**Table 1.**
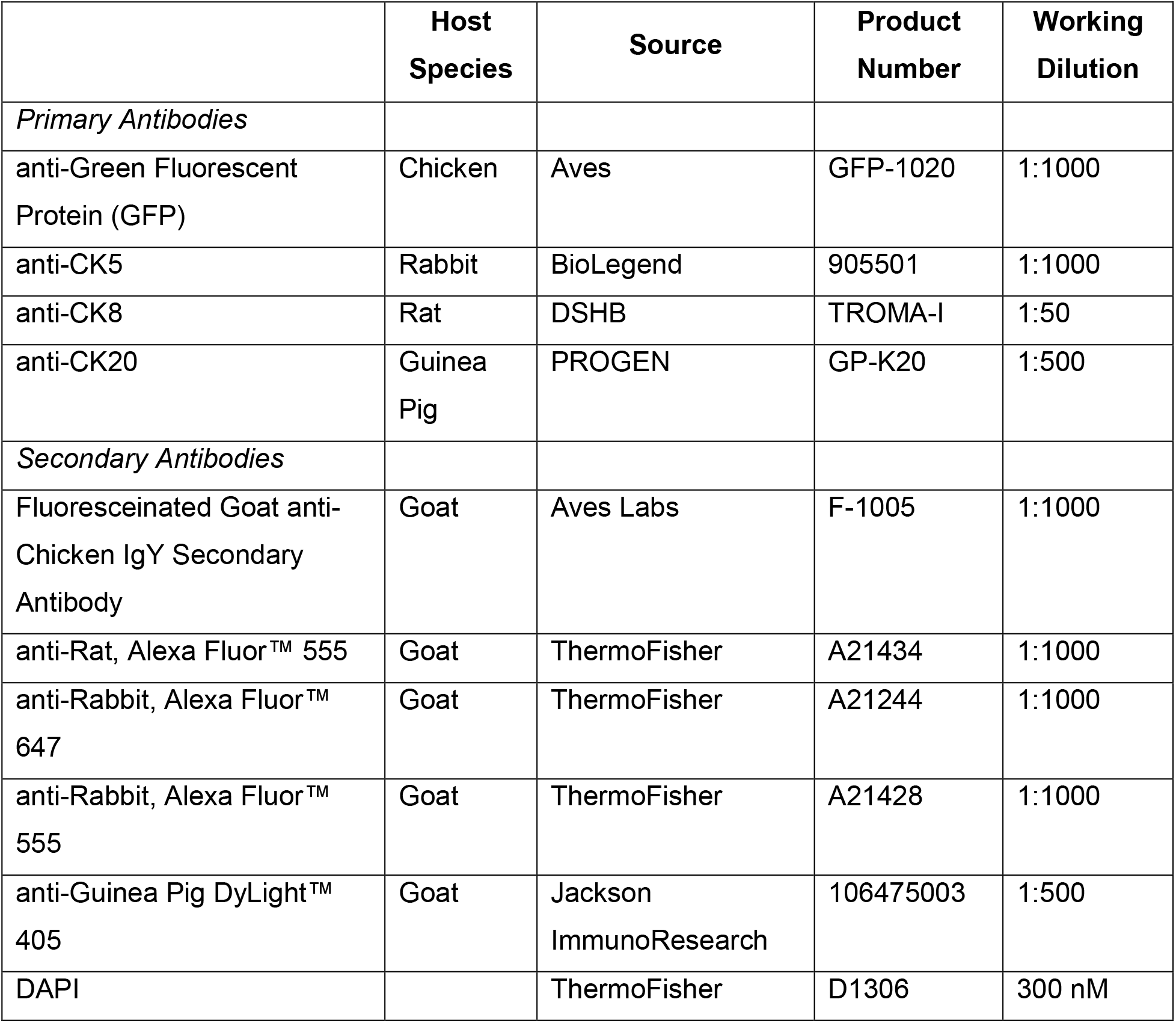
Antibodies used for immunohistochemistry

|  | **Host Species** | **Source** | **Product Number** | **Working Dilution** |
| --- | --- | --- | --- | --- |
| *Primary Antibodies* |  |  |  |  |
| anti-Green Fluorescent Protein (GFP) | Chicken | Aves | GFP-1020 | 1:1000 |
| anti-CK5 | Rabbit | BioLegend | 905501 | 1:1000 |
| anti-CK8 | Rat | DSHB | TROMA-I | 1:50 |
| anti-CK20 | Guinea Pig | PROGEN | GP-K20 | 1:500 |
| *Secondary Antibodies* |  |  |  |  |
| Fluoresceinated Goat anti-Chicken IgY Secondary Antibody | Goat | Aves Labs | F-1005 | 1:1000 |
| anti-Rat, Alexa Fluor™ 555 | Goat | ThermoFisher | A21434 | 1:1000 |
| anti-Rabbit, Alexa Fluor™ 647 | Goat | ThermoFisher | A21244 | 1:1000 |
| anti-Rabbit, Alexa Fluor™ 555 | Goat | ThermoFisher | A21428 | 1:1000 |
| anti-Guinea Pig DyLight™ 405 | Goat | Jackson ImmunoResearch | 106475003 | 1:500 |
| DAPI |  | ThermoFisher | D1306 | 300 nM |

### 3D Organoid Cell Culture

4-7 week old animals were sacrificed and an abdominal incision was made to locate the bladder^37^. The bladder was removed by one clean cut with sharp scissors at the bladder neck. The tissue was placed into a petri dish containing cold PBS (10010023, Gibco™). One end of the scissors was inserted into the base of the bladder and cut up the dome then opened to visualize the urothelium. Using blunt forceps, the urothelium was gently removed from the muscularis. The urothelium was washed in fresh, cold PBS and then placed in a 1.5 mL microcentrifuge tube containing: 200 μL 4.5 mg/mL Collagenase II (C6885, Sigma-Aldrich), 0.5 μL 10 mM Y-27632 (10005583, Cayman Chemical Company), 20 μL 10 mg/mL Elastase (E7885, Sigma-Aldrich), and 800 μL Advanced DMEM/F12+++ medium (see reagents and product numbers in **Table 2**). The urothelium was minced with sharp scissors and placed on a thermomixer (VWR Thermal Shake Touch, 89232-908, VWR) at 700 RPM for 2 hours at 37°C. The tissue digestion was gently vortexed every 15-30 minutes. After that, the tissue digestion was centrifuged at 750 x g for 5 minutes. The supernatant was aspirated, and the pellet was resuspended in 1 mL TrypLE™ Express Enzyme (12-605-010, Gibco™). The suspension was placed back on the thermomixer at 700 RPM for 30 minutes at 37°C. The suspension was centrifuged at 750 x g for 5 minutes, followed by aspiration of the supernatant. The pellet was resuspended in 1 mL Advanced DMEM/F12+++ media, and the number of cells was counted using a hemocytometer. The suspension was centrifuged at 750 x g for 5 minutes followed by aspiration of the supernatant. The pellet was resuspended in the appropriate amount of Cultrex Basement Membrane Extract (353300102, R&D Systems). 50 μL tabs were plated onto 6 well-plates (FB012927, Fisher Scientific) and incubated at 37°C for 30 minutes before 2 mL/well of Complete Growth Medium (see reagents and product numbers in **Table 3**) was added. Cultures were maintained at 37°C in 5% CO_2_ humidity-controlled incubators. Media was changed every other day and cells were passed at 70-80% confluency. Cells were maintained for up to 4 passages.

**Table 2.** Advanced DMEM/F12+++ Medium (per 500 mL)

| Reagent | Volume | Source | Product Number |
| --- | --- | --- | --- |
| Advanced DMEM/F12 | 500 mL | Gibco™ | 12634010 |
| GlutaMAX™ Supplement | 5 mL | Gibco™ | 35050061 |
| Penicillin/Streptomycin | 5 mL | Sigma-Aldrich | P4333 |
| HEPES | 5 mL | Gibco™ | 15630080 |

**Table 3.** Complete Growth Medium (per 50 mL)

| Reagent | Volume | Source | Product Number |
| --- | --- | --- | --- |
| B-27™ Supplement | 1 mL | Gibco™ | 17504044 |
| N-Acetyl-L-cysteine | 125 µL | Sigma-Aldrich | A9165 |
| Nicotinamide | 500 µL | Sigma-Aldrich | N0636 |
| mEGF | 2.5 µL | Gibco™ | PMG8041 |
| A-83-01 | 20 µL | Sigma-Aldrich | SML0788 |
| Recombinant Human Noggin | 100 µL | PeproTech | 120-10C |
| Recombinant Human R-Spondin-1 | 25 µL | PeproTech | 120-38 |
| Advanced DMEM/F12+++ Medium (Advanced DMEM/F12 supplemented with GlutaMAX™, Penicillin/Streptomycin, HEPES) | 44.853 mL | (See **Table 2**) | (See **Table 2**) |

### Immunocytochemistry

Once cells in culture reached 70-80% confluency, the organoids were collected and transferred to a 15 mL conical tube with equal parts Advanced DMEM/F12+++ medium. Organoids were then gently vortexed and placed on ice for 15-20 minutes. The cells were centrifuged at 750 x g for 5 minutes. The supernatant was removed, and the pellet was resuspended in 1 mL TrypLE™ Express Enzyme (12-605-010, Gibco™). Cells were incubated at 37°C in 5% CO_2_ humidity-controlled incubators for 10 minutes, gently vortexing every 5 minutes. 1-2 mL Advanced DMEM/F12+++ medium was added, and the cells were centrifuged at 750 x g for 5 min. The supernatant was removed, the pellet was resuspended in 1 mL Advanced DMEM/F12+++ medium, and the cells were counted using a hematocytometer. The appropriate amount of cell suspension was plated onto poly-d-lysine/collagen^38^ coated coverslips (∼20,000 cells/coverslip) and incubated at 37°C in 5% CO_2_ humidity-controlled incubators for 1-4 hours before each well was flooded with 1 mL CnT-Prime media (CNT-PR, ZenBio). Cultures were maintained at 37°C in 5% CO_2_ humidity-controlled incubators until fixation (48-72 hours).

Cells were fixed in 4% PFA for 10 minutes at room temperature followed by 3 washes in 1x PBS for 10 minutes each. Non-specific antibody binding was blocked with a blocking solution (3% normal goat serum, 0.3% Triton™ X-100, in 1x PBS) for 45 minutes at room temperature. Cells were incubated with primary antibodies (**Table 1**) for 1 hour at room temperature followed by 3 washes in 1x PBS for 10 minutes each. Thereafter, cells were incubated with secondary antibodies (**Table 1**) for 45 minutes at room temperature followed by 3 washes in 1x PBS for 10 minutes each. All antibodies were diluted in the blocking solution. Coverslips were mounted onto slides using VECTASHIELD Vibrance Antifade Mounting Media. All images were captured with the Keyence BZ-X Series all-in-one fluorescence microscope, BZ-X Viewer and, processed with Fiji ImageJ (1.53t, NIH).

### Patch Clamp Electrophysiology

Organoid cell cultures were split, as described in the previous section, onto poly-d-lysine/collagen-coated coverslips 24-48 hours before recording. These cells are attached to the coverslips as plaques with flat morphologies. Coverslips were transferred to the recording chamber with an extracellular buffer containing (in mM): 135 NaCl, 10 KCl, 10 HEPES, 10 Glucose, 1.2 MgCl_2_, and 1 CaCl_2_. The pipette solution contained (in mM): 145 KCl, 0.2 CaCl_2_, 1 MgCl_2_, 5 EGTA, and 10 HEPES. These solutions were previously used in urothelial patch-clamp recordings^19^. Currents were recorded at room temperature (∼22_o_C) with an Axopatch 200B patch-clamp amplifier connected to a Digidata 1440A data acquisition system (Molecular Devices). The holding potential was -60 mV, and the data were sampled at 2 kHz and filtered at 1 kHz using pClamp 10 software (Molecular Devices). Light stimulation was applied using LEDs and a multi-LED driver through the microscope optics (DC4100, Thor Labs). The intensity was measured using digital optical power and energy meter (PM100D; Thor Labs). Patch pipettes were pulled (PC-10, Narashige) from borosilicate glass tubes (TW150F-4, World Precision Instruments) and heat polished at the tip using a microforge (MF-200, World Precision Instruments) to give a resistance of 2–6 MΩ when filled with the pipette solution. Clampfit 10 (Molecular Devices), Excel (Microsoft Co), and Prism 9 (GraphPad Software) were used for the analysis of currents and preparing traces/figures. A Zeiss Axioscope was used to identify GFP-expressing cells and guide the pipette to the cell with a Sutter micromanipulator (MP-225).

### ATP Release Assay

Organoid Cell cultures were split onto black 24-well microplates (82426, Ibidi) 24-48 hours before the assay. Cells were washed 3 times with 300 μL HBSS+H. 200 μL of POM-1 (2689, Tocris) working stock^39^ was added to each well, and plates were incubated at 37°C for 30 minutes to allow for ATP degradation that may have been released during prior steps. Positive control wells were treated with 100 nM or 1 nM of the TRPV4 agonist, GSK1016790A (6433, Tocris), and incubated for 15 minutes at room temperature^32^. 200 μL from each positive control well was collected into cold microcentrifuge tubes and then put immediately onto the ice. Sample wells were then treated with 20 mW LED (M470F3, Thor Labs) connected to a fiber optic patch cable which was positioned ∼0.5 cm above the media surface, at 37°C for either 1, 5, or 10 minutes. 200 μL from each sample well was collected into cold microcentrifuge tubes and then put immediately onto the ice. Luminescence was recorded using CellTiter-Glo® 2.0 reagent (G9242, Promega) according to the manufacturer’s instructions using white, opaque 96 well-plates (3912, Corning) and a microplate reader (30-6006, Lmax II, Molecular Devices) equipped with SoftMax Pro 5.4. All samples were run in triplicate, each plate contained both positive and negative controls, and ATP (FLAAS, Sigma-Aldrich) standard curves were generated for each plate according to the Promega CellTiter-Glo® 2.0 protocol. The ATP standard curves calculated each sample’s ATP concentration using linear regression.

### *In vivo* Bladder Pressure Recordings

Mice were given a subcutaneous injection of urethane at 1.72 g/kg. After 1 hour, the mice were placed under 2% isoflurane through a nose cone. Once a negative pain response was detected using a toe pinch, an incision was made into the abdomen to reveal the bladder and a purse-string suture was made around the apex of the bladder. Using small spring scissors, the apex was punctured through to the bladder lumen and a fiber optic catheter was placed into the opening. The fiber optic catheter is custom built using optical fiber (FT200UMT, Thor Labs) running down polyethylene tubing (BB31695-PE/3, Scientific Commodities, Inc) which terminates as a Y-connector (1ZKE9, Grainger) where the optic goes through an epoxy sealed side and connects to the laser (473 DPSS Laser, BL472T8, Shanghai Laser & Optics Century Co., Ltd.). The fluid side connects to an inline pressure transducer (503067, World Precision Instruments) and a syringe pump (GT1976 Genie Touch, Kent Scientific Corporation). A solid-state pressure sensor (Scisense 1.2F Pressure catheter, TH-1211B-0018, Transonic) was inserted into the lumen, which was found to be more sensitive. The purse string suture was tightened around the fiber optic catheter and solid-state pressure sensor. After the catheter was secured, the skin and muscle incisions were closed with 5-0 suture (07-809-8813, Patterson Veterinary) and the isoflurane was lowered to zero.

### Continuous Flow Cystometry

1 hour after isoflurane cessation, the syringe pump was turned on at a flow rate of 0.04 mL/minute and a voiding pattern was established. Once regular contractions were established the experiment began. If a regular pattern could not be established the mouse was excluded from the experiment and further testing. Bladder pressure was measured using the inline pressure transducer connected to a Transbridge amplifier (TBM4, World Precision Instruments). The data was recorded using Micro 1401 (Cambridge Electronic Design) and Spike2 Software (V10; Cambridge Electronic Design).

### Light Intensity Pressure Assessments

One hour after the isoflurane was stopped, the syringe pump was started to demonstrate the bladder was capable of voiding upon filling. If no regular voiding pattern was generated, the experiment was terminated. The syringe pump was stopped, and the bladder was emptied manually. Then using the syringe pump, the bladder was filled with saline to about 40% of the volume needed to elicit a void. In this state, the different laser light intensities were tested. The inline pressure transducer (503067, World Precision Instruments) was used for these experiments. The data was recorded using Micro 1401 and Spike2 Software.

### PPADS experiments

After 1-hour post-isoflurane, the bladder was filled with saline to a state of half-capacity (about 5 mmHg). The bladder was stimulated with a 473 nm laser at 5 mW in 5-minute intervals. Once a baseline of 3 stimulations was established, saline was infused into the bladder, and the bladder was given 20 minutes to reach the same fill state as the baseline. Three stimulations were used to ensure that the administration of the liquid itself was not causing changes in response. This procedure was then repeated with the addition of the P2X receptor antagonist, PPADS (iso-PPADS tetrasodium salt, 0683, Tocris). Bladder pressure was recorded using the Scisense pressure transducer, and changes in bladder pressure were recorded using Micro 1401 and Spike2 Software.

### *Ex Vivo* Bladder Contraction Assay

The whole bladder preparation and recording of ex vivo bladder intravesical pressure measurements were performed as described previously^40^. Under brief exposure to 2% isoflurane, the mice were sacrificed by cervical dislocation. The urinary bladder and urethra were quickly excised and placed into a petri dish filled with physiological saline solution. The bladder was then freed of the connective tissues and the associated adipose. After that, the bladder was washed twice by inserting a polyethylene catheter through the remaining urethra, connected to a 23 G needle, and a syringe containing physiological saline solution. A fiber optic was fitted into a polyethylene catheter (as described above), and a Transonic pressure sensor was inserted into the bladder cavity through the urethra and sutured with silk thread to prevent urethral leakage. The prepared bladder was then transferred into a 10 mL organ bath containing warmed (37°C) physiological solutions (continuously carbonated with 95% O_2_ and 5% CO_2_), and physiological solutions were changed once the solution temperature went down to 30°C. The composition of the physiological saline solution (in mM) is: 119 NaCl, 4.7 KCl, 24 NaHCO_3_, 1.2 KH_2_PO_4_, 2.5 CaCl_2_, 1.2 MgSO_4_, and 11 Glucose with a pH = 7.3∼7.4. The catheter was connected to a syringe pump (GT1976 Genie Touch, Kent Scientific Corporation) to infuse the bladder with saline at a rate of 70-90 μL/minute, up to a final volume of 100-150 μL, and equilibrated for at least 1 hour before the experiment. The whole bladder contractile viability was assessed by application of 3 μM carbachol (L06674, Thermo Fisher Scientific) into the organ bath and washed 3 to 5 times with warm (37°C) physiological saline after achieving the maximum pressure. The pressure data was acquired with the Transonic Scisense pressure sensor and processed at a 1 kHz sampling rate using the Micro 1401 data acquisition system. The blue light (470 nm laser at 2-50 mW, 10-second continuous stimulation in 2–3-minute intervals) was applied to stimulate the urothelial ChR2.

### *In vivo* Electrophysiological Recording of Pelvic Nerve Afferents

Mice were anesthetized with isoflurane and a suprapubic bladder catheter with a fiber optic was implanted. A midline incision in the skin and muscle was made over the lower abdomen to expose the dome of the bladder. The bladder catheter with fiber optic and Transonic pressure sensor was implanted and secured into the dome of the bladder with a purse-string suture. The catheter was made from the aforementioned polyethylene tubing with flared ends. The catheter was connected via a 3−way valve to a syringe filled with saline for bladder distension. After the catheter was placed, the muscle and skin were sutured, and the animal was gently placed in the prone position. A dorsal incision was made between the lower lumbar vertebrae and the tail head to surgically isolate the pelvic nerve in the ischiorectal fossa (as described in^41^). The pelvic nerve was isolated, cut centrally, and placed on 36-gauge platinum-iridium hook electrodes bathed with mineral oil. Data was collected with Power 1401 (Cambridge Electronic Design). Raw nerve activity was integrated using Rectigy and integrated script from CED (version updated 06/16, https://ced.co.uk/downloads/scriptspkanal, Cambridge Electronic Design) with a 0.5-second integration time. Post-hoc data were analyzed and graphed with Spike2 and GraphPad Prism 9.

### Data Analysis

Data analysis and graphical construction were performed using GraphPad Prism 9 software. Quantitative results are represented as mean ± SEM. We used one-way ANOVA, two-way ANOVA, and paired student’s t-tests depending on the number of variables and groups for comparison. Each data point on the graphs represents a different animal or cell culture. Figures designed and produced using Affinity Designer V1 (Serif).

## Results

### Characterizing ChR2 Expression and Function in Urothelial Cells

We crossed UPK2-cre mice with ChR2-GFP (Ai32 mice) to create the UPK2-ChR2 mouse model. We found that ChR2-GFP was expressed in intermediate and some umbrella cells (**Figure 1**), as previously shown by others using the UPK2 promoter cre mouse^13,36^. ChR2-GFP expression was maintained in isolated urothelial cell culture (**Figure 1C**). We next characterized the cellular responses to optogenetic stimulation of urothelial cells in culture (**Figures 2A and B**). Stimulation of these cells with blue light resulted in membrane depolarization that was not present when cells were stimulated with amber light (**Figure 2C**). These currents increased with increased light intensity (**Figure 2D**). As mechanical and chemical stimulation can evoke ATP release from urothelial cells^22,42^, we evaluated whether optogenetic activation of these cells could initiate ATP release. We found that blue light stimulation of these cells resulted in ATP release in similar amounts to stimulation with 100 nM of the TRPV4 agonist, GSK1016790A (**Figure 2E**). These results suggest that the increase in intracellular cations (including Na^+^ and Ca^2+^) evoked by ChR2 activation is sufficient to initiate ATP release.

**Figure 1.**
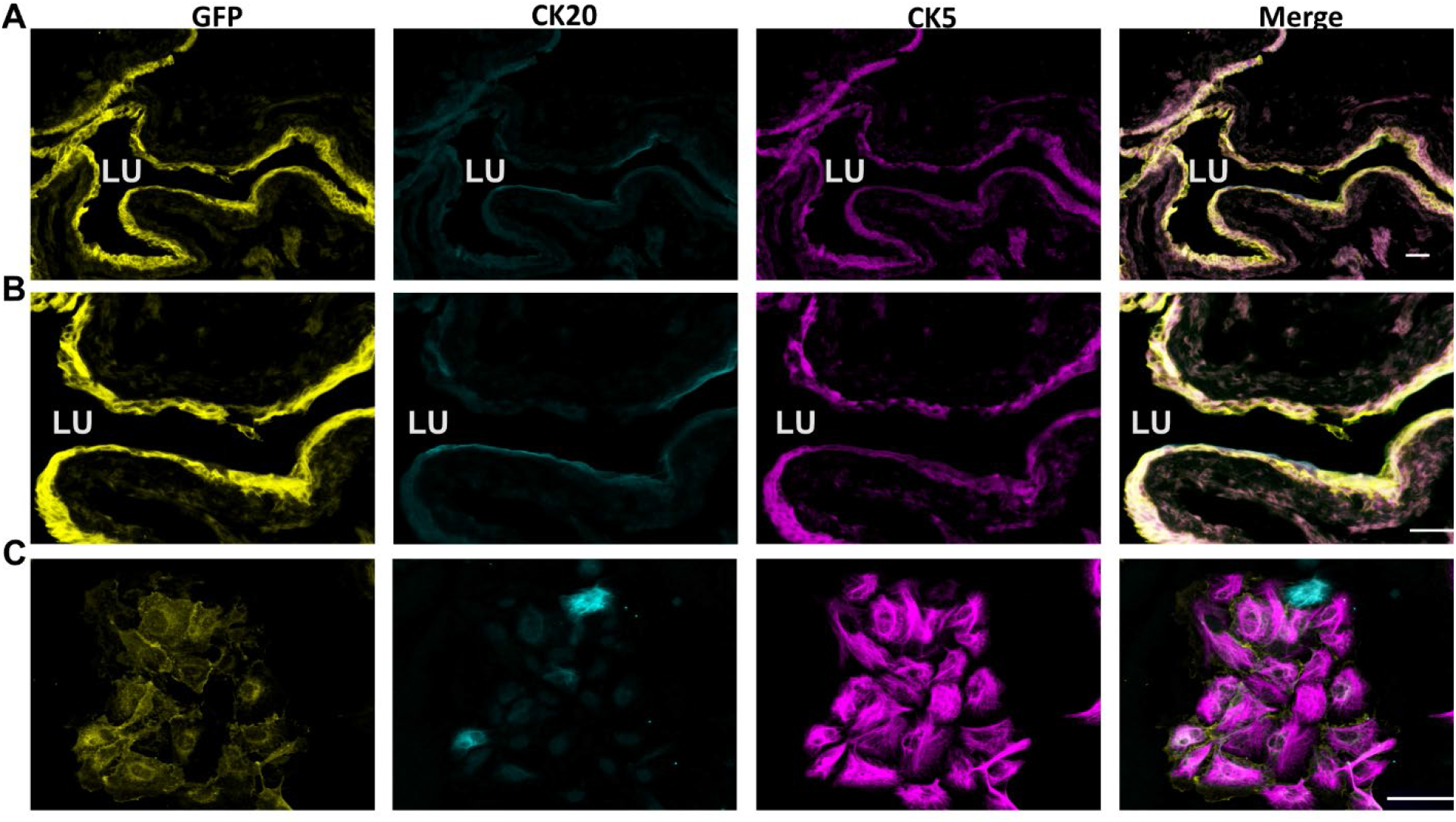
UPK2-ChR2 mice express ChR2-eGFP in urothelial cells. Mice were generated by crossing UKP2-cre with Ai32 (Flox-ChR2), referred to as UPK2-ChR2. Bladder cryosection (**A, B**) and isolated urothelial cells in culture (**C**) with ChR2-eGFP expression (yellow), co-stained with urothelial markers CK20 (cyan) and CK5 (magenta). LU, lumen; CK20, cytokeratin 20; CK5, cytokeratin 5; GFP, green fluorescent protein; Scale bars; 50 μm.

**Figure 2.**
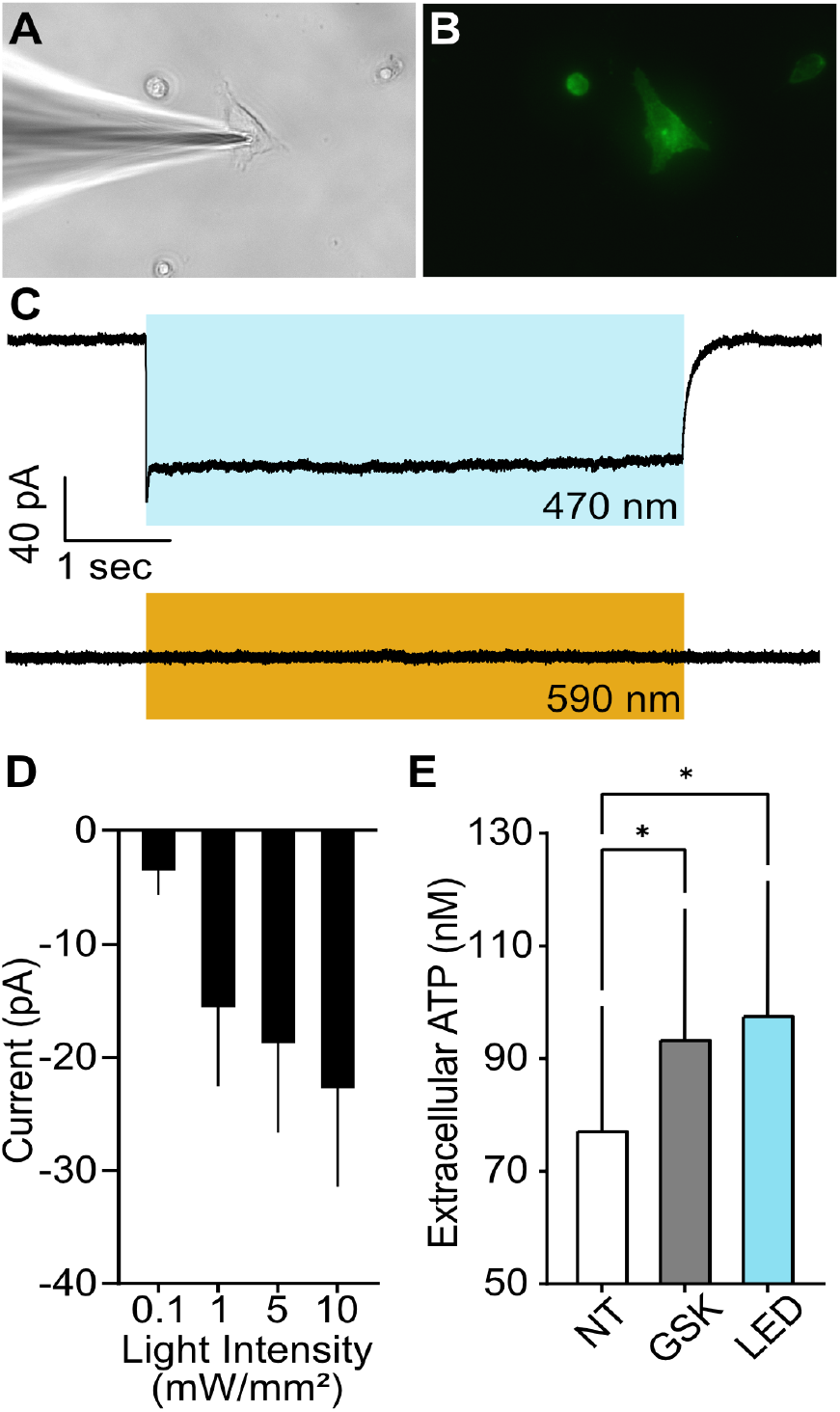
Effects of optogenetic stimulation on membrane potential and ATP release. Urothelial cells cultured from UPK2-ChR2 mice (**A**) were identified by GFP expression (**B**) and patched. Blue light evoked an inward current, and stimulation with amber light did not affect membrane currents (**C**). (**D**) The optically evoked currents increased with the increased blue light intensities. (**E**) Quantification of ATP release from UPK2-ChR2 cultured cells. Cells were treated with media containing 100 nM GSK1016790A (GSK), a TRPV4 agonist, or blue light stimulation (LED; 20 mW). n= 3 different cultures. *p<0.05 compared to no treatment (NT) controls. Repeated Measures One-way ANOVA, Dunnett’s Multiple Comparison (NT vs GSK p= 0.011 and NT vs LED p=0.023).

### Effects of Optogenetic Stimulation on Bladder Pressure

We next wanted to see the effect of optogenetic-mediated urothelial activation on bladder pressure. Using a catheter containing a fiber optic we performed continuous flow cystometry and then measured the pressure changes in response to light stimulation. 10 mW light stimulation evoked bladder contractions of similar pressure to cystometric voids (**Figures 3A and B**). This light-initiated response was repeatable outside the normal contraction pattern. In mice where continuous infusion was stopped, and bladders were filled to approximately 40% voiding threshold, we evaluated the relationship between light intensity and change in intraluminal pressure. We found a positive correlation between light intensity and light-evoked contraction magnitude (**Figures 3C and D**). This correlation supports the hypothesis that urothelial stimulation can influence the bladder contraction state. Increasing the intensity presumably activates more ChR2 receptors in the urothelium and initiates larger contractions (**Figure 3E**). Importantly, blue light stimulation of the bladder from cre-negative mice demonstrated no contractions confirming that ChR2 expression is required for the blue light to evoke bladder contractions.

**Figure 3.**
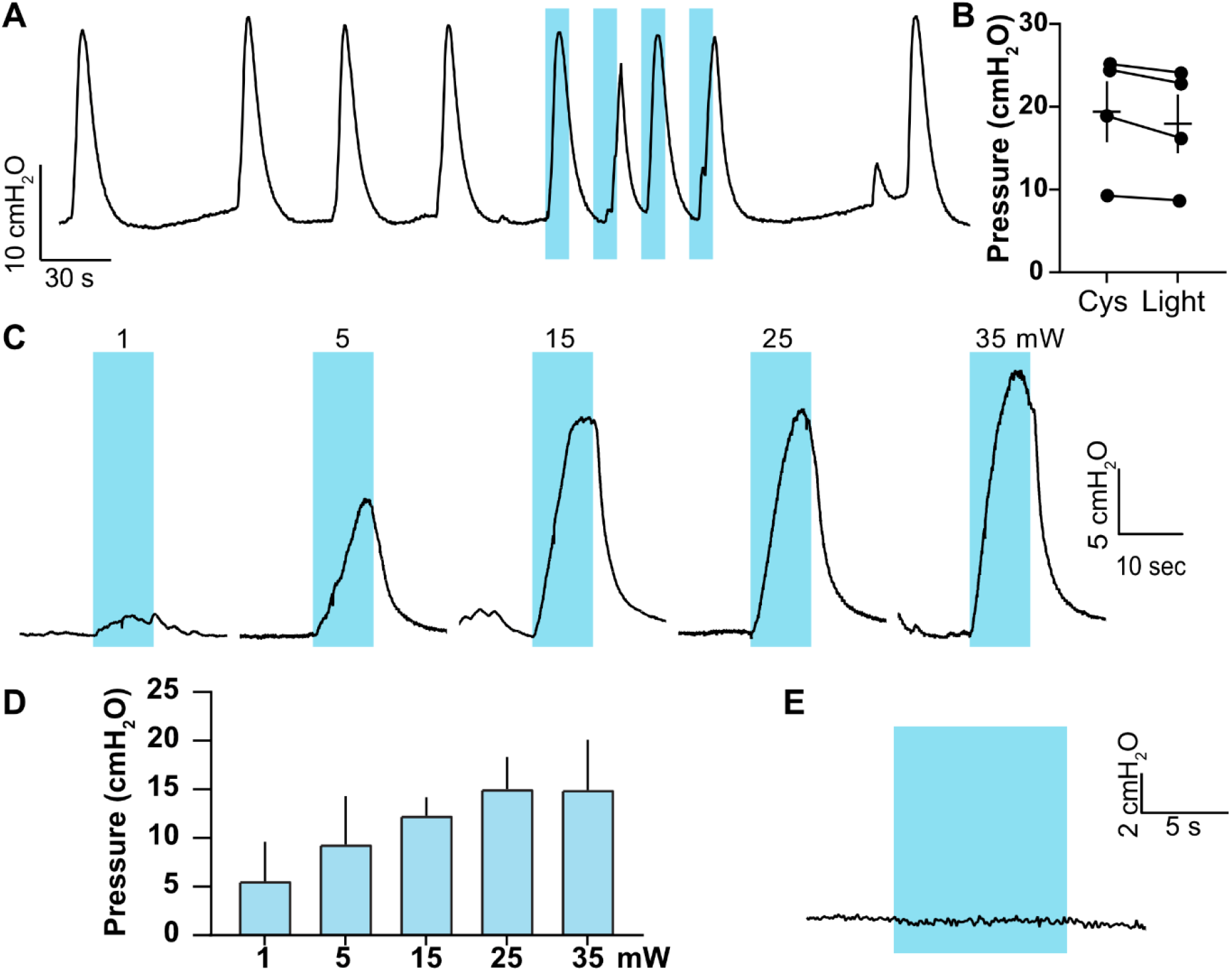
Optogenetic urothelial stimulation in UPK2-ChR2 mice is sufficient to evoke bladder contractions. (**A**) Repeated 10-second optical stimulation of the bladder urothelium evoked bladder contractions outside the normal contraction pattern during continuous flow cystometry (10 mW). (**B**) These ChR2-induced contractions (Light; 18.04 ± 3.549 cmH2O) were of a similar magnitude to cystometric voids (Cys;19.48 ± 3.686 cmH2O). (**C** and **D**) In a half-full bladder, increasing the intensity of light stimulation increases the magnitude of pressure response in UPK2-ChR2 mice. (**E**) Blue light stimulation in UPK2-ChR2 Cre negative mice could not evoke contractions, demonstrating the effect of blue light was related to the expression of ChR2.

Next, we wanted to evaluate if the contraction evoked by the optogenetic stimulation of the urothelial cells was a local signaling event or if it required afferent signaling to the central nervous system to produce a reflex contraction. In our first experiment, we evaluated the urothelial-evoked contractions before and after the unilateral severing of the pelvic nerve. This resulted in decreased optically evoked bladder contractions (**Figure 4A**), suggesting that these contractions are partly mediated by the pelvic nerve afferent and centrally mediated reflex voiding.

**Figure 4.**
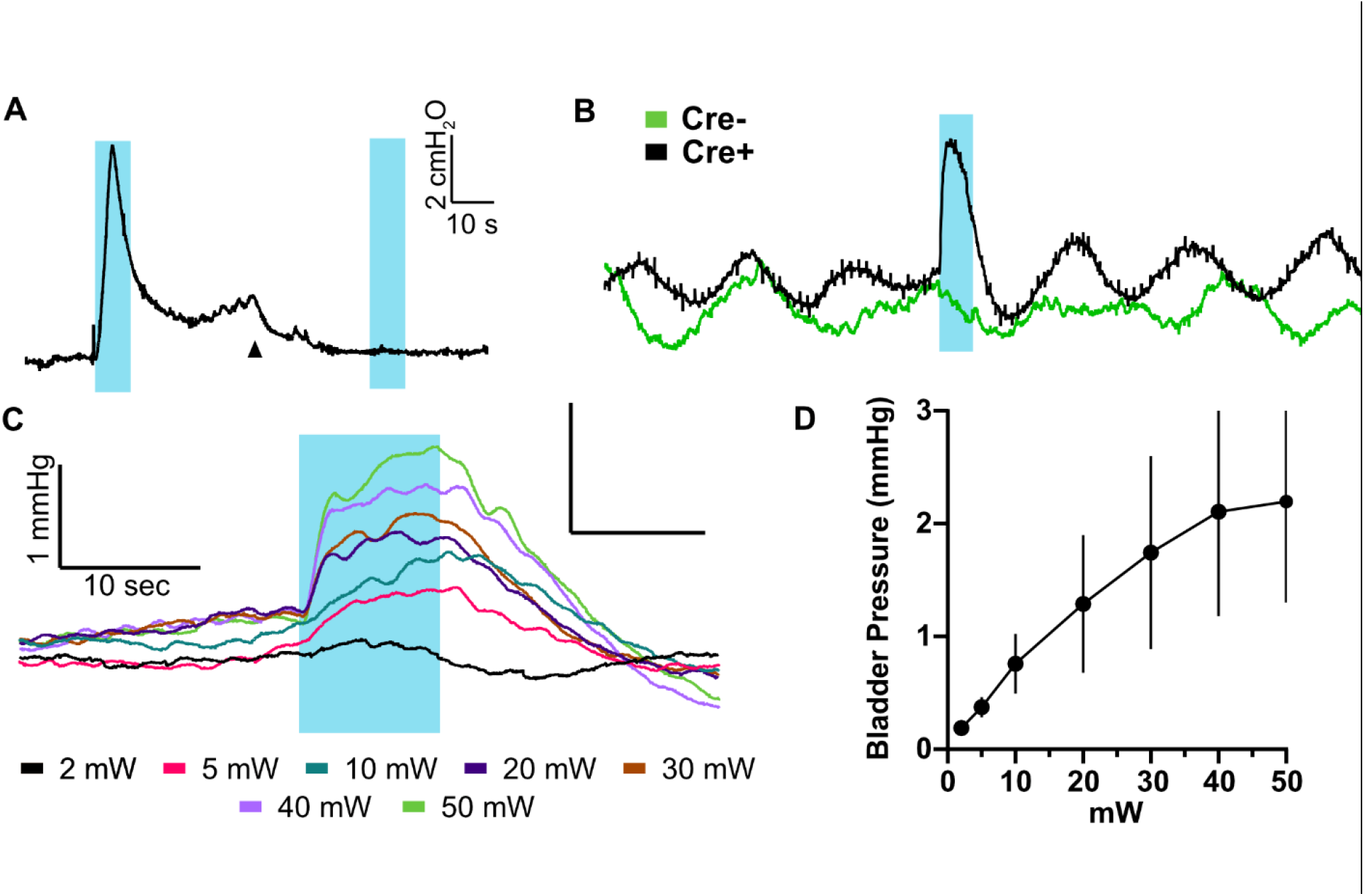
ChR2-mediated simulation of urothelial cells includes locally mediated bladder contractions. (**A**) Unilateral transection of the pelvic nerve (black arrow) reduced optogenetically evoked contractions *in vivo*. (**B**) Optogenetic stimulation (20 mW) of urothelial cells *ex vivo* resulted in small bladder contractions distinct from background contractions in bladders from Cre+ mice (black trace) but not in Cre-mice (green trace). (**C**) Example traces from *ex vivo* urothelial stimulation at increasing light intensities and (**D**) quantification of pressure response to increasing light intensity (n=9; Mean ± SEM).

We then performed an *ex vivo* bladder experiment to test if optogenetic urothelial stimulation could evoke bladder contractions without central circuits. We found that blue light stimulation could repeatably produce changes in intraluminal pressure greater than the normal phasic contractions (**Figures 4B - D**). Together, these results suggest that urothelial stimulation evoked increases in bladder pressure that involves a local signaling event in addition to signaling and communication with the central nervous system.

### Optogenetic Stimulation of Urothelial Cells Evokes Pelvic Sensory Neuron Activity

Urothelial cells are expected to communicate directly or indirectly with terminations of sensory neurons, so we wanted to determine if optogenetic urothelial stimulation can evoke activity in centrally projecting pelvic afferents. Using an *in vivo* electrophysiology preparation, we recorded whole nerve (multi-unit) activity from the distal end of the transected severed distal pelvic nerve in UPK2-ChR2 mice. With this approach, we could evaluate the sensory neuron responses without interference from descending efferent projections. Optogenetic stimulation of urothelial cells from bladders of UPK2-ChR2 mice evoked an increase in sensory neuron activity (**Figure 5**). Light pulses (1 second on, 1 second off) correlated with increases in nerve activity (**Figure 5A**). The delay from light stimulation to increased nerve activity was relatively quick (24.7 ± 4.27 ms), and peak response was reached in 1.05 ± 0.27 sec (**Figures 5B - F**). Longer stimulations resulted in the same increase in activity that corresponded with an increase in bladder pressure (**Figure 5D**). Once the bladder pressure reached its peak, the nerve activity decreased despite the continued light stimulation (**Figures 5D and E**).

**Figure 5.**
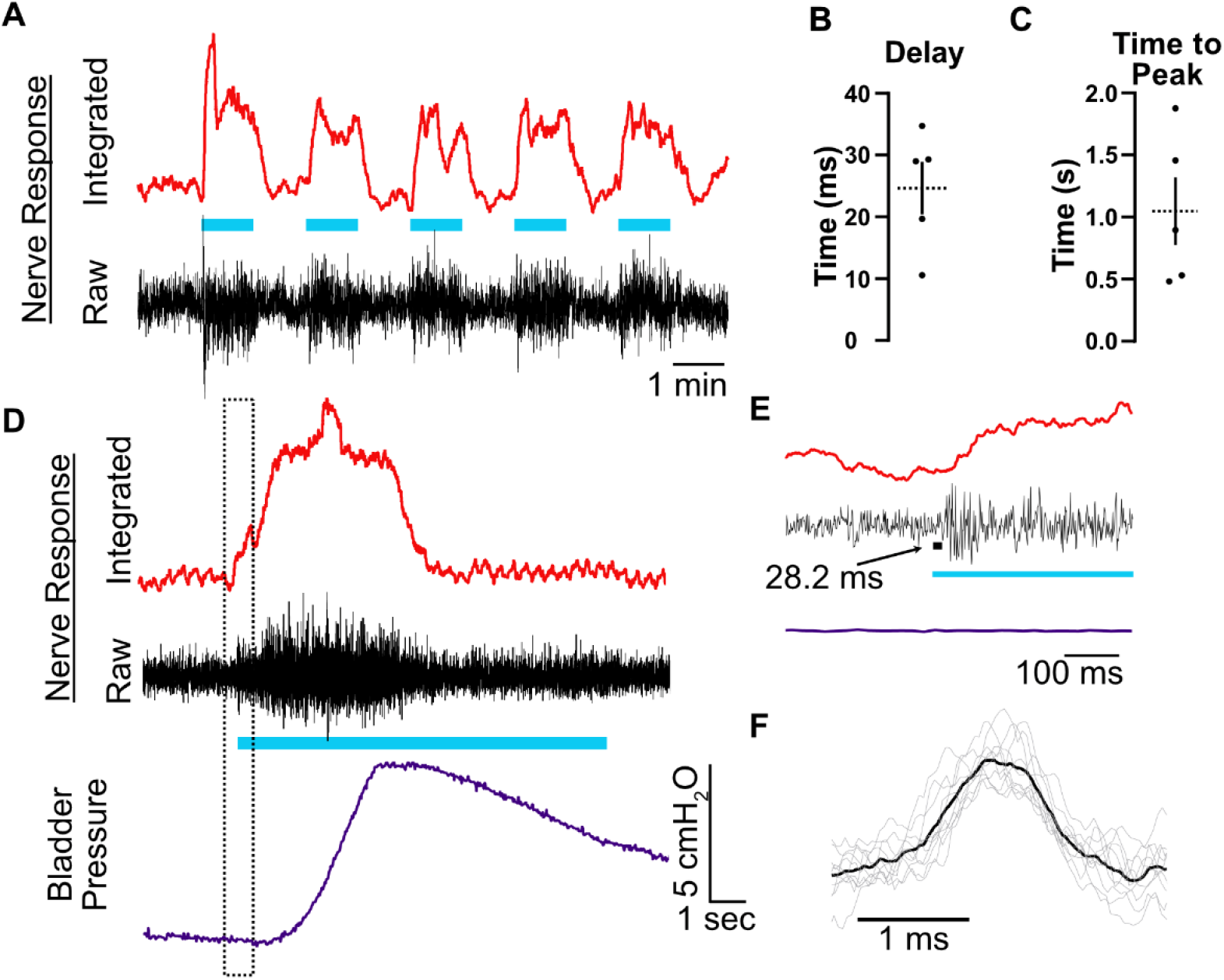
Urothelial stimulation evokes sensory pelvic nerve activity. (**A**) A representative trace of distal cut-end pelvic nerve activity (black trace) in response to optogenetic urothelial stimulation (blue bars; 1 sec on 1 sec off). The integrated nerve activity (red trace) can delineate the increase in nerve activity during light pulses. (**B**) The increase in nerve activity was initiated at an average of 24.7 ± 4.27 ms after the optical stimulation, and (**C**) reached its peak activity in 1.05 ± 0.27 sec following the optical stimulation (n=5). (**D**) Longer stimulation (Blue bar, 10 seconds) resulted in an increase in nerve activity followed by an increase in bladder pressure. Once bladder pressure peaked, nerve activity decreased. (**E**) Close-up of panel D in the dotted box demonstrating the 28.2 ms latency between onset of light stimulation and the onset of pelvic nerve afferent firing. (**F**) An average action potential waveform (the black line is the average of grey waveforms) of spikes (n=10) during optical stimulation. Note that the action potential duration is consistent with those of unmyelinated C-fibers as is the expected conduction latency of ∼15-20 ms.

### Role of Purinergic Signaling in Urothelial Evoked Bladder Contractions and Afferent Nerve Activity

ATP and purinergic signaling play an important role in normal bladder function. To test the role of purinergic signaling in urothelial optogenetic-mediated bladder contractions, we stimulated the urothelial cells before and after the instillation of the pan-P2X receptor inhibitor, PPADS, into the bladder. We found that 4 mM PPADS could significantly reduce the maximum pressure elicited by optogenetic stimulation of bladders from UPK2-ChR2 mice (**Figures 6A and B**). This effect could be reversed by washing out the PPADS (**Figure 6C**). We then recorded pelvic nerve activity evoked by optogenetic urothelial stimulation before and after intravesical delivery of 1mM PPADS (**Figure 6D**). We observed a significant decrease in pelvic nerve activity after the delivery of PPADS as quantified by the percent change area under the curve in the integrated nerve response (**Figure 6E**). PPADs reduced pelvic nerve activity to 31.22 ± 12.32% of baseline response.

**Figure 6.**
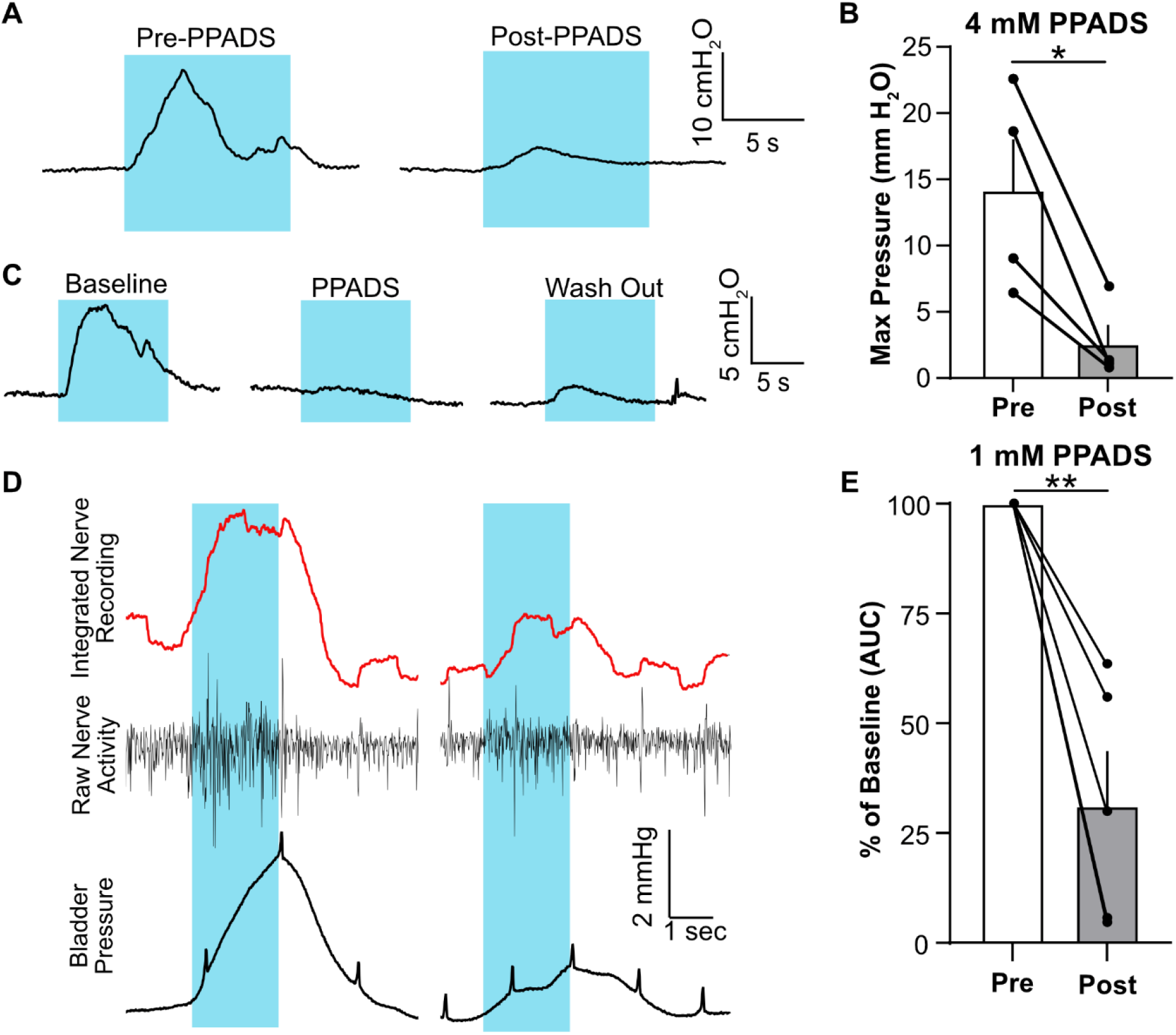
Inhibition of P2X receptors reduces bladder contractions and sensory pelvic nerve firing evoked by urothelial stimulation. Example traces (**A**) and quantification (**B**) of ChR2 evoked contractions before and during treatment with intravesical PPADS (4 mM). PPADS could reduce the max peak of the contraction. *p=0.0285 Paired T-test (n=4). (**C**) Example traces demonstrate that after washout of PPADS, contractions could be rescued. Blue bars indicate when light stimulation is delivered. (**D-E**) Non-specific inhibition of P2X receptors with 1 mM PPADS reduced the whole pelvic nerve activity associated with optogenetic activation of urothelial cells. Small black spikes are the respiration influence on the abdominal cavity and bladder. Paired T-test **p=0.0051 (n=5). Light blue bars indicate light stimulation.

Furthermore, we wanted to test the importance of P2X receptor signaling in the local urothelial stimulation-induced contractions. Here we observed that 1 mM PPADS reduced and almost eliminated the local optogenetic-induced contractions, suggesting that P2X receptor signaling is essential for local urothelial-induced contractions (**Figures 7A and B**).

**Figure 7.**
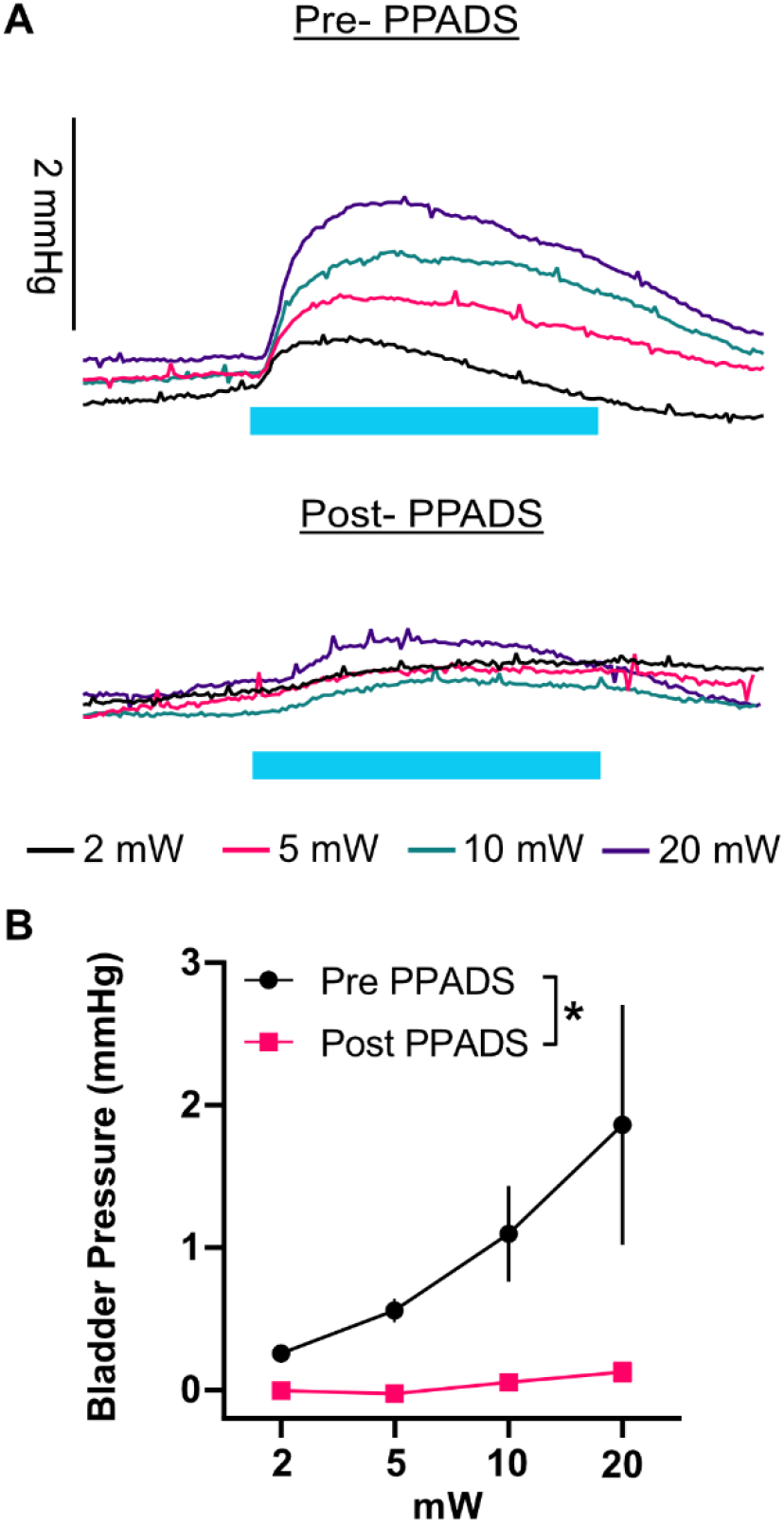
Inhibition of P2X receptors significantly reduces local urothelial-mediated contractions. (**A**) Representative traces *ex vivo* bladder pressure from UPK2-ChR2 blue light simulation (blue bar). (**B**) Quantification of the percent reduction of maximum contraction pressure before and after 1 mM PPADS. (n=7, mean ± SEM, p=0.0375 comparing treatment groups, 2-way ANOVA).

## Discussion

### Sensory Function of Urothelial Cells

Urothelial cells respond to sensory signals in the bladder, including mechanical and chemical stimuli, and release different chemical mediators. However, it has been a challenge to study the role of this signaling to sensory neurons, as they express many of the same sensory receptors as neuronal endings innervating the urothelial layer. Here we have demonstrated a new optogenetic-based method to investigate the communication between urothelial cells and other cell types in the bladder that allows for the specific activation of urothelial cells without directly affecting other bladder cell types. We found that optogenetic stimulation of urothelial cells results in contractions of similar magnitude as cytometric contractions (**Figure 3**). Further optogenetic stimulation of urothelial cells quickly (∼20 ms) increased the pelvic nerve C-fiber-like neuronal activity (**Figure 5**), supporting the idea of a fast communication mechanism between urothelial cells and sensory neurons.

This work aligns the foundation of studies supporting a sensory role for urothelial cells, by showing that direct and specific stimulation of urothelial cells was sufficient to increase sensory nerve activity and evoke bladder contractions. Removing afferent signaling to the CNS and efferent control of the detrusor by severing the pelvic nerve or excising the bladder (*ex vivo*) from the mouse reduced the magnitude of contraction. These results along with the nerve recording data, suggest that sensory signaling plays an important role in the observed optogenetic-mediated urothelial bladder contractions.

### Locally Mediated Contractions

In our *ex vivo* bladder pressure recordings from UPK2-ChR2 mice, we observed that optogenetic-mediated stimulation of the urothelium evoked an increase in intraluminal pressure above the phasic contractions (**Figure 4**). These results indicate that in the absence of centrally mediated reflex contractions urothelial stimulation can still induce bladder contractions. Two potential mechanisms for this action are contraction of the lamina propria and urothelial layers and signaling through the lamina propria mediating a detrusor contraction.

Isolated urothelium/lamina propria spontaneously develop phasic contractile activity and non-phasic contractions can be evoked by electrical stimulation, neurokinin A, neuropeptide Y, ATP, or acetylcholine^43–46^. In this study, we observed that inhibition of P2X receptors with PPADS could nearly abolish optically evoked contractions, indicating ATP’s importance in urothelial-stimulated local contractions. Electrical stimulation evoked contraction of the urothelial/lamina propria layer, which presumably activates efferent and afferent nerve endings, is not blocked by purinergic receptor inhibition^46^ indicating that the urothelial stimulated contractions likely involve a different mechanism. Further studies are needed to determine what P2X receptor expressing cells are responsible for this action, as many cells within the bladder wall express P2X receptors.

Alternatively, optogenetic urothelial stimulation could induce the release factors that influence detrusor activity in evoking a contraction. The lamina propria lies between the urothelium and the detrusor and may form structural and physiological links between urothelial cells, sensory nerves, and detrusor muscle cells^47,48^. Therefore, an alternative hypothesis is that optogenetic urothelial activation influences detrusor activity mediating the contractions. For example, the greater amplitude of bladder spontaneous contraction in intact full-thickness bladder wall strips than the bladder strips with dissected lamina propria and urothelium suggested that lamina propria has a positive modulatory influence on the detrusor muscle contraction^49^.

### Urothelial Evoked Pelvic Nerve Afferent Activity

It has long been hypothesized that urothelial cells transmit sensory signals to sensory nerve afferents due to the proximity and expression of many receptors able to respond to urothelial released factors. Here we demonstrate that optogenetic activation of urothelial cells results in quick and robust activation of sensory neurons of the pelvic nerve. While we were able to record smaller bundles of fibers with a low background to distinguish single fibers, we were able to show that these spikes are in the range of 1-3 ms in duration, consistent with unmyelinated C-fibers. Further, these neurons have a distinct audio signature of C-fibers, and not A-delta myelinated sensory fibers that also innervate the bladder. Future studies will focus on recordings from smaller teased branches will allow for the identification of single spikes to identify these neurons further. Additional recordings in disease models will indicate if the populations responding to optogenetic stimulation will be modified under disease conditions.

### The Role of P2X Receptor in Urothelial Evoked Bladder Contractions

Inhibition of P2X receptors resulted in a decrease in pressure and nerve activity evoked by urothelial stimulation. We also demonstrated that optogenetic urothelial stimulation resulted in a local bladder-mediated effect on bladder pressure dependent on P2X receptor signaling. This evidence supports the importance of purinergic signaling in this communication mechanism.

Purinergic receptor signaling plays an important role in the micturition reflex and bladder nociception^42^. P2X receptors are found on nerve fibers innervating the bladder as well as small sensory neurons^34,50^. Patch clamp recordings from DRG neurons innervating the urethra or distal colonic mucosa and traveling through the pelvic nerve exhibit depolarizing currents elicited by ATP and acetylcholine^51^. Early studies have found that P2X_3_ and P2X_2/3_ knock-out mice experience reduced nociceptive behaviors after ATP administration as well as bladder hyporeflexia^34^. However, a more recent study has shown that the P2X_3_, P2X_2/3_ knock-out mice, and PPADS-treated wild-type mice did not have altered bladder function under normal physiological conditions, although they did see attenuated bladder hyperactivity under pathological conditions^52^. In our study, we showed that optically stimulated urothelial cells could communicate with sensory neurons to initiate bladder contractions, and the administration of a P2X receptor antagonist, PPADS, attenuates nerve activity and contraction. Based on the literature and our data, we can say that purinergic signaling plays a fundamental role in urothelial cell communication, but its specific function still needs to be elucidated.

This work demonstrates that purinergic signaling is important for urothelial-driven bladder contractions, but we cannot confirm that ATP released by urothelial cells is acting on sensory neurons directly. As urothelial cells express, purinergic receptors ATP and P2X receptors may be an important autocrine signaling mechanism to amplify sensory responses with the urothelium. Further, inhibiting P2X receptors with PPADS leads to a ∼70% reduction in nerve activity, suggesting that there is another component involved in the sensory nerve response to urothelial stimulation. ATP can also evoke the release of acetylcholine from the urothelium and act on sensory afferents to evoke reflex contractions^53–55^. Further studies are needed to see if acetylcholine is downstream or in parallel with P2X receptor signaling in urothelial stimulation-evoked contractions.

### Study limitations

While this new technique has further supported the hypothesis that urothelial cells communicate with sensory neurons, we cannot determine from this study if it is direct communication. The short delay between the optical stimulation of the urothelium and the increase in nerve activity suggests direct communication. However, the signaling could also go through a secondary relay through another cell, like interstitial cells^32,56,57^. Additionally, while we demonstrated that it does evoke ATP release (**Figure 2**), ChR2 is an exogenous channel that is not normally present in urothelial cells, and thus the downstream signaling initiated by its activation is unknown and likely differs from endogenous cation channels like TRPV4 or P2X_2_. This is a critical point to consider when interpreting these results. New opsins are being engineered to mimic endogenous receptor activity; future studies could utilize these to link at specific downstream signaling cascades^58^. Finally, the physiological effects of bladder contractions were observed under anesthesia. Future studies must evaluate the effect of urothelial-specific stimulation on non-anesthetized cystometric and non-cystometric voiding to confirm that urothelial stimulation can evoke independent voiding.

## Conclusions

In conclusion, we have demonstrated that urothelial stimulation alone in the absence of other stimuli is sufficient to evoke bladder contractions. We have established a technique to optically stimulate urothelial cells and study physiological outputs, including bladder cystometry function and pelvic nerve sensory afferent activity. The results from this work leave many unanswered questions but with a new approach to address cellular communication in the bladder. Like the influx of optogenetics techniques into the brain, these techniques will allow us to understand better the cellular communication concerning bladder physiology and pathophysiology. Importantly aberrant release of signaling molecules, like ATP, have been proposed as critical components of IC/BPS and OAB. Using these optogenetic or chemogenetic techniques to modulate urothelial cells specifically under disease models will hopefully better describe a role for this cell type communication in disease. Further expanded application of different types of opsins (G-protein coupled receptor, transcriptional regulators) will allow new questions to be answered.

## References

1. Grundy L, Erickson A, Brierley SM. Visceral Pain. Annu Rev Physiol. 2019;81(1):261–284. doi:10.1146/annurev-physiol-020518-114525

2. Birder LA, Wolf-Johnston AS, Chib MK, Buffington CA, Roppolo JR, Hanna-Mitchell AT. Beyond neurons: Involvement of urothelial and glial cells in bladder function. Neurourol Urodyn. 2010;29(1):88–96. doi:10.1002/nau.20747

3. Elbadawi A, Light JK. Distinctive Ultrastructural Pathology of Nonulcerative Interstitial Cystitis. Urol Int. 1996;56:137–162. doi:https://doi.org/10.1159/000282832

4. Tomaszewski JE, Landis JR, Russack V, et al. Biopsy features are associated with primary symptoms in interstitial cystitis: results from the interstitial cystitis database study. Urology. 2001;57(6):67–81.

5. Keay SK, Birder LA, Chai TC. Evidence for Bladder Urothelial Pathophysiology in Functional Bladder Disorders. BioMed Res Int. 2014;2014:15. doi:10.1155/2014/865463

6. Hurst RE, Greenwood-Van Meerveld B, Wisniewski AB, et al. Increased bladder permeability in interstitial cystitis/painful bladder syndrome. Transl Androl Urol. 2015;4(5). doi:10.3978/j.issn.2223-4683.2015.10.03

7. Lai MC, Kuo YC, Kuo HC. Intravesical hyaluronic acid for interstitial cystitis/painful bladder syndrome: A comparative randomized assessment of different regimens. Int J Urol. 2013;20:203–207. doi:10.1111/j.1442-2042.2012.03135.x

8. Kim JC, Park EY, Hong SH, Seo SI, Park YH, Hwang TK. Changes of urinary nerve growth factor and prostaglandins in male patients with overactive bladder symptom. Int J Urol. 2005;12(10):875–880. doi:10.1111/j.1442-2042.2005.01140.x

9. Kim JC, Park EY, Seo SI, Park YH, Hwang TK. Nerve Growth Factor and Prostaglandins in the Urine of Female Patients With Overactive Bladder. J Urol. 2006;175(5):1773–1776. doi:10.1016/S0022-5347(05)00992-4

10. Suh YS, Ko KJ, Kim TH, et al. Potential Biomarkers for Diagnosis of Overactive Bladder Patients: Urinary Nerve Growth Factor, Prostaglandin E2, and Adenosine Triphosphate. 2017;21:171–177. doi:10.5213/inj.1732728.364

11. Sun Y, Chai TC. Augmented extracellular ATP signaling in bladder urothelial cells from patients with interstitial cystitis. Am J Physiol-Cell Physiol. 2006;290(1):C27–C34. doi:10.1152/ajpcell.00552.2004

12. Marshall KL, Saade D, Ghitani N, et al. PIEZO2 in sensory neurons and urothelial cells coordinates urination. Nature. 2020;(588):290–295. doi:10.1038/s41586-020-2830-7

13. Dalghi MG, Carattino MD, Apodaca G. Functional roles for PIEZO1 and PIEZO2 in urothelial mechanotransduction and lower urinary tract interoception. JCI Insight. 2021;6(19). doi:10.1172/jci.insight.152984

14. Miyamoto T, Mochizuki T, Nakagomi H, et al. Functional Role for Piezo1 in Stretch-evoked Ca2+ Influx and ATP Release in Urothelial Cell Cultures. J Biol Chem. 2014;289(23):16565–16575. doi:10.1074/jbc.M113.528638

15. Kullmann FA, Shah MA, Birder LA, de Groat WC. Functional TRP and ASIC-like channels in cultured urothelial cells from the rat. Am J Physiol-Ren Physiol. 2009;296(4):F892–F901. doi:10.1152/ajprenal.90718.2008

16. Corrow K, Girard BM, Vizzard MA. Expression and response of acid-sensing ion channels in urinary bladder to cyclophosphamide-induced cystitis. Am J Physiol-Ren Physiol. 2010;298(5):F1130–F1139. doi:10.1152/ajprenal.00618.2009

17. Birder LA, Nakamura Y, Nealen ML, et al. Altered urinary bladder function in mice lacking the vanilloid receptor TRPV1. Nat Neurosci. 2002;5:856–860. doi:10.1038/nn902

18. Birder LA, Kullmann FA, Lee H, et al. Activation of Urothelial Transient Receptor Potential Vanilloid 4 by 4α-Phorbol 12,13-Didecanoate Contributes to Altered Bladder Reflexes in the Rat. J Pharmacol Exp Ther. 2007;323(1):227–235. doi:10.1124/jpet.107.125435

19. Li M, Sun Y, Simard JM, Chai TC. Increased transient receptor potential vanilloid type 1 (TRPV1) signaling in idiopathic overactive bladder urothelial cells. Neurourol Urodyn. 2011;30(4):606–611. doi:10.1002/nau.21045

20. Merrill L, Girard BM, May V, Vizzard MA. Transcriptional and Translational Plasticity in Rodent Urinary Bladder TRP Channels with Urinary Bladder Inflammation, Bladder Dysfunction, or Postnatal Maturation. J Mol Neurosci. 2012;48:744–756. doi:10.1007/s12031-012-9867-5

21. Alm P, Alumets J, Brodin E, et al. Peptidergic (Substance P) Nerves in the Genito-Urinary Tract. Neuroscience. 1978;3:419–425. doi:0306.4522/78/0501-041980

22. Ferguson DR, Kennedy I, Burton TJ. ATP is released from rabbit urinary bladder epithelial cells byhydrostatic pressure changes — a possible sensory mechanism? J Physiol. 1997;505(2):503–511. doi:10.1111/j.1469-7793.1997.503bb.x

23. Wang ECY, Lee JM, Ruiz WG, et al. ATP and purinergic receptor-dependent membrane traffic in bladder umbrella cells. J Clin Invest. 2005;115(9):2412–2422. doi:10.1172/JCI24086

24. Lewis SA, Lewis JR. Kinetics of urothelial ATP release. Am J Physiol-Ren Physiol. 2006;291(2):F332–F340. doi:10.1152/ajprenal.00340.2005

25. Lips KS, Wunsch J, Zarghooni S, et al. Acetylcholine and Molecular Components of its Synthesis and Release Machinery in the Urothelium. Eur Urol. 2007;51(4):1042–1053. doi:10.1016/j.eururo.2006.10.028

26. Beckel JM, Kanai A, Birder LA. Expression of functional nicotinic acetylcholine receptors in rat urinary bladder epithelial cells. Am J Physiol-Ren Physiol. 2009;290(1):F103–F110. doi:10.1152/ajprenal.00098.2005

27. Birder LA, Apodaca G, De Groat WC, Kanai AJ. Adrenergic- and capsaicin-evoked nitric oxide release from urothelium and afferent nerves in urinary bladder. Am J Physiol-Ren Physiol. 1998;275(2):F226–F229. doi:10.1152/ajprenal.1998.275.2.F226

28. Birder LA, Nealen ML, Kiss S, et al. B-Adrenoceptor Agonists Stimulate Endothelial Nitric Oxide Synthase in Rat Urinary Bladder Urothelial Cells. J Neurosci. 2002;22(18):8063–8070. doi:10.1523/JNEUROSCI.22-18-08063.2002

29. Dixon JS, Gilpin CJ. Presumptive sensory axons of the human urinary bladder: a fine structural study. J Anat. 1987;151:199–207.

30. Dixon JS, Jen PYP, Gosling JA. Immunohistochemical characteristics of human paraganglion cells and sensory corpuscles associated with the urinary bladder. A developmental study in the male fetus, neonate and infant. J Anat. 1998;192:407–415. doi:10.1046/j.1469-7580.1998.19230407.x

31. Birder L, Andersson KE. Urothelial signaling. Physiol Rev. 2013;93(2):653–680. doi:10.1152/physrev.00030.2012

32. Merrill L, Gonzalez EJ, Girard BM, Vizzard MA. Receptors, channels, and signaling in the urothelial sensory system in the bladder. Nat Rev Urol. 2016;(13):193–204. doi:10.1038/nrurol.2016.13

33. Vlaskovska M, Kasakov L, Rong W, et al. P2X3 knock-out mice reveal a major sensory role for urothelially released ATP. J Neurosci. 2001;21(15):5670–5677.

34. Cockayne DA, Hamilton SG, Zhu QM, et al. Urinary bladder hyporeflexia and reduced pain-related behaviour in P2X3-deficient mice. Nature. 2000;407(6807):1011–1015. doi:10.1038/35039519

35. Madisen L, Mao T, Koch H, et al. A toolbox of Cre-dependent optogenetic transgenic mice for light-induced activation and silencing. Nat Neurosci. 2012;15:793–802. doi:10.1038/nn.3078

36. Ayala de la Pena F, Kanasaki K, Kanasaki M, Tangirala N, Maeda G, Kalluri R. Loss of p53 and acquisition of angiogenic microRNA profile are insufficient to facilitate progression of bladder urothelial carcinoma in situ to invasive carcinoma. J Biol Chem. 2011;286(23):20778–20787. doi:10.1074/jbc.M110.198069

37. Halstead AM, Kapadia CD, Bolzenius J, et al. Bladder-cancer-associated mutations in RXRA activate peroxisome proliferator-activated receptors to drive urothelial proliferation. eLife. 2017;6:e30862. doi:10.7554/eLife.30862

38. Everaerts W, Vriens J, Owsianik G, et al. Functional characterization of transient receptor potential channels in mouse urothelial cells. Am J Physiol Renal Physiol. 2010;298(3):F692–F701. doi:10.1152/ajprenal.00599.2009

39. Durnin L, Moreland N, Lees A, Mutafova-Yambolieva VN. A commonly-used ecto-ATPase inhibitor, ARL-67156, blocks degradation of ADP more than the degradation of ATP in murine colon. Neurogastroenterol Motil. 2016;28(9):1370–1381. doi:10.1111/nmo.12836

40. Park JH, Hong JK, Jang JY, et al. Optogenetic Modulation of Urinary Bladder Contraction for Lower Urinary Tract Dysfunction. Sci Rep. 2017;7:40872. doi:10.1038/srep40872

41. Hubscher CH, Johnson RD. Responses of medullary reticular formation neurons to input from the male genitalia. J Neurophysiol. 1996;76(4):2474–2482. doi:10.1152/jn.1996.76.4.2474

42. Moro C, Uchiyama J, Chess-Williams R. Urothelial/lamina propria spontaneous activity and the role of M3 muscarinic receptors in mediating rate responses to stretch and carbachol. Urology. 2011;78(6):1442.e9-15. doi:10.1016/j.urology.2011.08.039

43. Sadananda P, Chess-Williams R, Burcher E. Contractile properties of the pig bladder mucosa in response to neurokinin A: a role for myofibroblasts? Br J Pharmacol. 2008;153(7):1465–1473. doi:10.1038/bjp.2008.29

44. Mattiasson A, Andersson KE, Sjogren C. Contractant and Relaxant Properties of the Female Rabbit Urethral Submucosa. J Urol. 1985;133(2):304–310. doi:10.1016/S0022-5347(17)48928-2

45. Moro C, Chess-Williams R. Non-adrenergic, non-cholinergic, non-purinergic contractions of the urothelium/lamina propria of the pig bladder. Auton Autacoid Pharmacol. 2012;32(3-4):53–59. doi:10.1111/aap.12000

46. McCloskey KD. Interstitial Cells of Cajal in the Urinary Tract. In: Urinary Tract. Vol 2011. Handbook of Experimental Pharmacology. Springer, Berlin, Heidelberg; 2011:233–254. https://link.springer.com/chapter/10.1007/978-3-642-16499-6_11#citeas

47. Ikeda Y, Kanai A. Urotheliogenic modulation of intrinsic activity in spinal cord-transected rat bladders: role of mucosal muscarinic receptors. Am J Physiol-Ren Physiol. 2008;295(2):F454–F461. doi:10.1152/ajprenal.90315.2008

48. Andersson KE, McCloskey KD. Lamina propria: The functional center of the bladder? Neurourol Urodyn. 2014;33(1):9–16. doi:10.1002/nau.22465

49. Andersson KE. Purinergic signalling in the urinary bladder. Auton Neurosci. 2015;191:78–81. doi:10.1016/j.autneu.2015.04.012

50. Tempest HV, Dixon AK, Turner WH, Elneil S, Sellers LA, Ferguson DR. P2X2 and P2X3 receptor expression in human bladder urothelium and changes in interstitial cystitis. BJU Int. 2004;93(9):1344–1348. doi:10.1111/j.1464-410X.2004.04858.x

51. Rau KK, Petruska JC, Cooper BY, Johnson RD. Distinct subclassification of DRG neurons innervating the distal colon and glans penis/distal urethra based on the electrophysiological current signature. J Neurophysiol. 2014;112(6):1392–1408. doi:10.1152/jn.00560.2013

52. Takezawa K, Kondo M, Kiuchi H, et al. Authentic role of ATP signaling in micturition reflex. Sci Rep. 2016;6:19585. doi:10.1038/srep19585

53. Stenqvist J, Winder M, Carlsson T, Aronsson P, Tobin G. Urothelial acetylcholine involvement in ATP-induced contractile responses of the rat urinary bladder. Eur J Pharmacol. 2017;809:253–260. doi:10.1016/j.ejphar.2017.05.023

54. Stenqvist J, Carlsson T, Winder M, Aronsson P. Effects of caveolae depletion and urothelial denudation on purinergic and cholinergic signaling in healthy and cyclophosphamide-induced cystitis in the rat bladder. Auton Neurosci. 2018;213:60–70. doi:10.1016/j.autneu.2018.06.001

55. Stenqvist J, Aronsson P, Carlsson T, Winder M, Tobin G. In vivo paracrine effects of ATP-induced urothelial acetylcholine in the rat urinary bladder. Auton Neurosci. 2020;227. doi:10.1016/j.autneu.2020.102689

56. Li Y, Xue L, Miao Q, et al. Expression and electrophysiological characteristics of P2X3 receptors in interstitial cells of Cajal in rats with partial bladder outlet obstruction. BJU Int. 2012;111(5):843–851. doi:10.1111/j.1464-410X.2012.11408.x

57. Sui GP, Fry CH. Electrical Characteristics of Suburothelial Cells Isolated From the Human Bladder. J Urol. 2004;171(2):938–943. doi:10.1097/01.ju.0000108120.28291.eb

58. Emiliani V, Entcheva E, Hedrich R, et al. Optogenetics for light control of biological systems. Nat Rev Methods Primer. 2022;2(55). doi:10.1038/s43586-022-00136-4

